# DeMoP: A Language-Model-Guided Mixture-of-Experts Framework for Cancer Prognosis

**DOI:** 10.64898/2026.08.24.746579

**Authors:** Chen Tang, Lei Yu, Qiwei Li, Lin Xu

## Abstract

Integrating heterogeneous clinical and molecular data for cancer prognosis remains challenging because their dimensionality, semantics and distributions differ across patients and cohorts. Here we present DeMoP, a language-model-guided mixture-of-experts framework that serializes structured patient profiles as natural-language sequences and learns adaptive prognostic representations from clinical variables, copy-number alterations, and gene descriptions. DeMoP combines a fine-tuned DeBERTa-v3-large encoder, attention-based token pooling, and a residual mixture-of-experts prediction head. In held-out tests from two independent pan-cancer cohorts, GENIE (63,090 patients) and TCGA (4,123 patients), DeMoP outperformed the conventional machine-learning and deep-learning baselines evaluated, achieving AUROCs of 0.939 and 0.805 and class-1 F1 scores of 0.72 in both cohorts. A GENIE-trained model transferred directly to TCGA with an overall class-1 F1 score of 0.62. Gene-level ablations recovered established cancer-associated genes and highlighted less-studied candidates. DeMoP provides a unified approach to heterogeneous biomedical data integration, cross-cohort outcome prediction, and model interpretation.

## Introduction

Accurate cancer prognosis is essential for treatment selection, clinical trial stratification, and long-term disease management. However, predicting patient outcomes remains challenging because clinical and molecular data are intrinsically heterogeneous^1,2^. Clinical variables such as age, sex, and tumor stage are relatively low-dimensional and interpretable, whereas molecular profiles, including somatic mutations, copy-number alterations, gene expression, and epigenetic features, are high-dimensional and complex^3,4^.

Traditional statistical and machine-learning approaches often rely on domain-specific feature engineering to integrate these heterogeneous modalities. Such approaches are labor-intensive and may miss nonlinear, higher-order interactions between clinical and genomic factors associated with patient outcomes^5,6^. Recent advances in natural language processing (NLP) and pretrained language models, including encoder-based models such as RoBERTa^7^ and DeBERTa^8^, offer an alternative strategy for modeling biomedical data. By representing structured patient information as natural-language-like sequences, transformer architectures can learn cross-modal relationships within a shared representation space^9^. In this setting, heterogeneous clinical and molecular attributes are serialized into token sequences and processed by self-attention, enabling flexible integration without bespoke feature engineering.

General-purpose biomedical large language models such as BioGPT^10^ and Med-PaLM^11^ have shown strong performance in medical question answering, clinical reasoning, and text understanding. However, these models are primarily designed for unstructured clinical text and are not optimized for structured multi-omic or tabular patient data, limiting their applicability to integrative prognostic modeling. In parallel, transformer-based approaches for structured and tabular data, including TabPFN^12^, TabTransformer^13^, and FT-Transformer^14^, have improved performance over conventional machine-learning methods by modeling feature interactions with self-attention. Although effective for tabular prediction, these approaches usually assume fixed feature schemas and do not readily accommodate heterogeneous biomedical modalities or leverage large-scale pretrained language representations.

More recently, LLM-inspired approaches have been extended to biological sequences and single-cell data. Models such as scGPT^15^, CELLama^16^, and sciLaMA17 adapt transformer architectures to encode gene-expression profiles and molecular features as tokenized representations, enabling downstream tasks such as cell-type annotation and perturbation modeling. Despite their promise, these approaches primarily focus on cell-level problems and have not been systematically evaluated for patient-level outcome prediction across heterogeneous real-world cohorts.

To address these challenges, we present DeMoP, a language-model-guided framework for cancer prognosis. DeMoP combines a fine-tuned DeBERTa-v3-large encoder (24 transformer layers; 304 million backbone parameters and 131 million embedding parameters)^8,18^ with concatenated representations from its final four hidden layers, an attention-based pooling module, and a ResNet-based mixture-of-experts head. DeBERTa’s disentangled attention and relative positional encoding support fine-grained modeling of structured patient representations, whereas the MoE head enhances the model’s adaptability through input-dependent specialization across heterogeneous patient profiles. Unlike prior approaches restricted to single cancer types or more homogeneous datasets, DeMoP is designed for cross-cohort generalization and integration of heterogeneous clinical and molecular data. We evaluate DeMoP in two independent pan-cancer cohorts, AACR Project GENIE^19^ and the TCGA database^20^, and benchmark it against conventional machine-learning methods and deep-learning architectures.

## Results

### Overview of DeMoP

DeMoP encodes patient-level clinical characteristics (e.g., sex and age) and gene-level information (e.g., copy-number alterations and brief gene descriptions) as unified structured natural-language representations (**Fig. 1**), enabling heterogeneous clinical and molecular data to be processed within a common framework.

**Figure 1.**
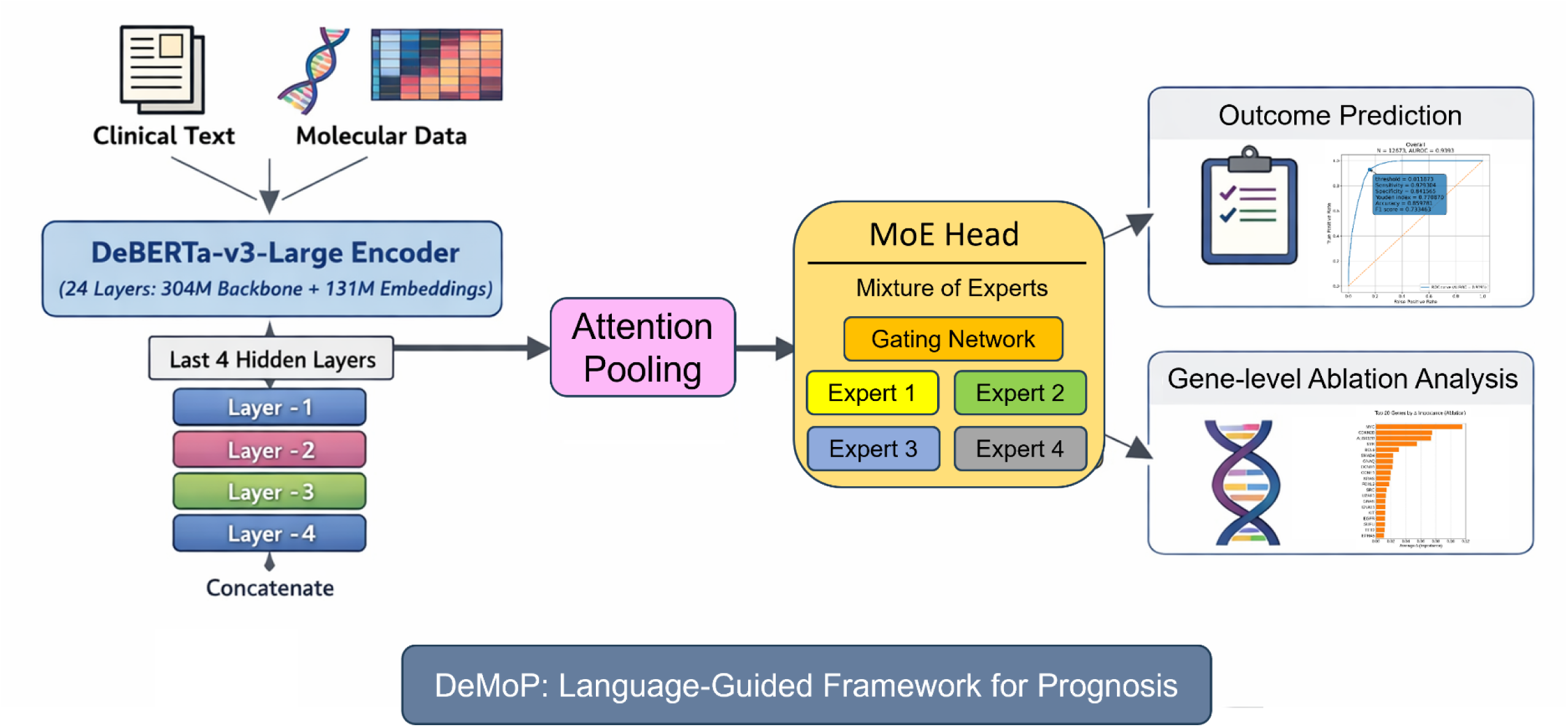
Schematic overview of the DeMoP framework. Patient-level clinical and molecular features are serialized into structured natural-language representations and encoded by a fine-tuned DeBERTa-v3-large. Representations from the final four hidden layers are concatenated and processed by an attention-based pooling module and a ResNet-based mixture-of-experts prediction head for pan-cancer outcome prediction and gene-level ablation analysis.

These representations are passed through a fine-tuned DeBERTa-v3-large encoder (24 transformer layers; 304 million backbone parameters and 131 million embedding parameters). Concatenated representations from its final four hidden layers are then used to aggregate complementary contextual information across semantic depths, conceptually inspired by YOLOv2-style feature aggregation^21^. Together, these components form the clinical and molecular embedding module of DeMoP.

Patient embeddings are then passed to a mixture-of-experts (MoE) head, in which a gating network assigns input-dependent weights to multiple specialized expert subnetworks based on each patient’s embedding. This design enables adaptive modeling of heterogeneous patient representations through complementary nonlinear patterns that may not be fully captured by a single prediction head. Together with the contextual patient embeddings, the MoE head constitutes the prediction module of DeMoP (**Methods: Mixture-of-experts head**).

To evaluate DeMoP on real-world clinical and molecular data, we conducted three complementary computational experiments: (1) training and evaluation using the AACR Project GENIE cohort (63,090 patients), (2) training and evaluation using the TCGA cohort (4,123 patients), and (3) a transfer analysis in which a model trained on the larger AACR Project GENIE cohort was evaluated on the smaller TCGA cohort, which included overlapping cancer types. Across these settings, we compared DeMoP with several conventional machine-learning and deep-learning approaches.

### DeMoP achieves strong predictive performance in the GENIE cohort

As a large real-world oncology cohort, AACR Project GENIE provides a testbed for evaluating performance across 84 diverse cancer types (**Methods: Dataset collection and preprocessing**). For the GENIE cohort, samples from patients who were alive but had less than three years of follow-up were excluded. This filtering reduced the number of samples from 130,124 to 64,681. Samples with zero expression across all genes in the 3-year mortality group were then removed. After quality-control filtering, 63,090 samples remained in the final GENIE cohort. We trained DeMoP using 80% of the GENIE cohort and evaluated it on the remaining 20% held-out internal test set. **Fig. 2A** compares DeMoP with six state-of-the-art benchmark methods on the GENIE test set after application of the predefined survival-based inclusion criteria. For 3-year survival prediction, DeMoP achieved a class-1 F1 score of 0.72, outperforming baseline models using default parameter settings: logistic regression (0.51), random forest (0.58), XGBoost (0.66), multilayer perceptron (0.57), CNN (0.63), and LSTM (0.59), all evaluated using the same training and test sets as DeMoP. The performance margin over both conventional classifiers and other deep-learning architectures is consistent with the integration of gene-level copy-number alteration profiles and diverse clinical annotations. For large patient cohorts aggregated across hospitals or medical centers, this integrative approach may help model heterogeneity in real-world clinical data.

**Figure 2.**
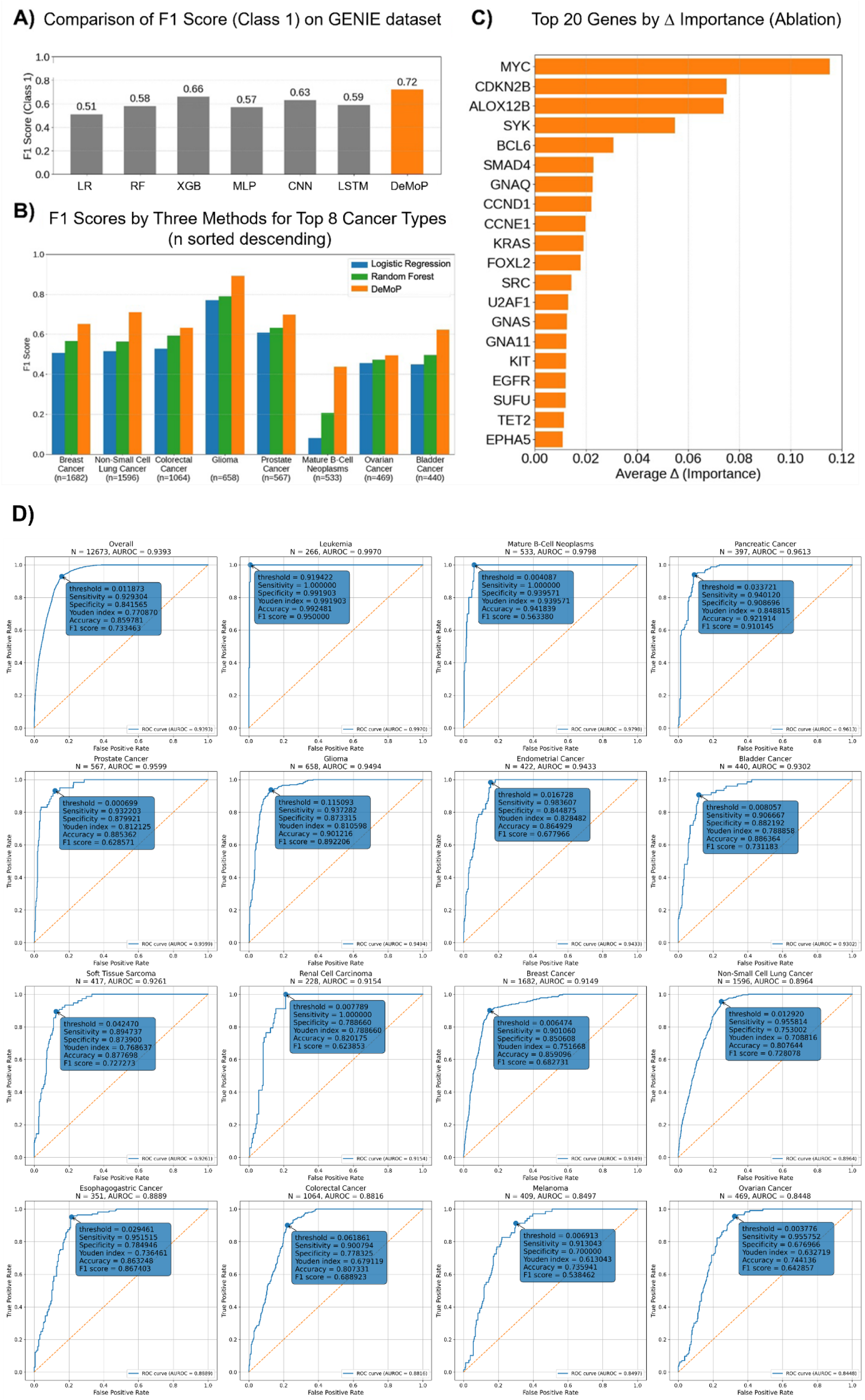
DeMoP performance in the AACR Project GENIE cohort. (A) Class-1 F1 scores on the held-out GENIE test set for DeMoP and all benchmark methods. (B) Class-1 F1 score comparisons across eight cancer types, with DeMoP outperforming random forest in common solid tumors and more than doubling the logistic regression score in mature B-cell neoplasms. (C) Top 20 genes ranked by the mean signed change in prediction confidence (Δ) in the ablation analysis, with MYC, CDKN2B, ALOX12B, and SYK among the highest-ranked genes. (D) Overall and cancer-specific AUROC performance of DeMoP on the GENIE test set.

**Fig. 2B** shows class-1 F1 scores for the eight cancer types with the largest test-set sample sizes in the GENIE cohort. In breast cancer (n = 1,682), non-small cell lung cancer (n = 1,596), and colorectal cancer (n = 1,064), DeMoP achieved higher class-1 F1 scores than random forest (0.65 versus 0.57, 0.71 versus 0.56, and 0.63 versus 0.59, respectively). In glioma (n = 658), DeMoP achieved a class-1 F1 score of 0.89, compared with 0.79 for random forest. In prostate cancer (n = 567), DeMoP achieved a class-1 F1 score of 0.70, compared with 0.63 for random forest and 0.61 for logistic regression. In mature B-cell neoplasms (n = 533), DeMoP more than doubled the class- 1 F1 score of logistic regression (0.44 versus 0.08). DeMoP also achieved modestly higher class- 1 F1 scores than random forest in ovarian cancer (0.50 versus 0.47) and bladder cancer (0.62 versus 0.50), indicating performance gains across solid and hematological malignancies.

To examine gene-level model dependence, we performed a systematic gene-level ablation analysis using DeMoP (**Methods: Ablation study)**. For each gene, we removed the corresponding gene-level segment, including its copy-number alteration data and associated brief descriptive text, from each patient description, passed the modified input through the fine-tuned model, and measured the resulting change in predicted probability for the model’s original predicted class relative to the full input. We defined each gene’s importance score as the mean reduction in prediction confidence upon omission of its gene-level information across all patients in the cohort (**Methods: Ablation study, Eq. 21**); a larger positive score indicates stronger model dependence on the encoded copy-number and descriptive information for that gene. The 20 genes with the highest mean importance scores are shown in **Fig. 2C**. MYC had the highest importance score (0.12), consistent with its central role in cell proliferation across cancer types^22^. CDKN2B (0.07) also ranked highly, in line with its role in cell-cycle control^23,24^. ALOX12B (0.07), a lipoxygenase implicated in ferroptosis and inflammation^25^, ranked among the top features but relatively less studied, suggesting that the model may capture previously underexplored microenvironmental dependencies. SYK (0.05), a kinase involved in B-cell receptor signaling^26^, ranked fourth and is consistent with the strong performance of DeMoP in mature B-cell neoplasms. Other highly ranked genes included BCL6 (0.03), a key regulator of germinal-center formation and B-cell differentiation^27^, and SMAD4 (0.02), a critical mediator of TGF-β signaling with established tumor-suppressor functions^28^.

At the cohort level, DeMoP achieved an overall AUROC of 0.939, with sensitivity of 0.772, specificity of 0.901, a Youden index of 0.673, accuracy of 0.874, and a weighted F1 score of 0.718. We next examined cancer-specific discrimination among tumor types with more than 200 samples in the test set (**Fig. 2D**). Among 15 cancer types, the highest AUROC was observed in leukemia (n = 266, AUROC = 0.997), followed by mature B-cell neoplasms (n = 533, AUROC = 0.980), pancreatic cancer (n = 397, AUROC = 0.961), prostate cancer (n = 567, AUROC = 0.960), and glioma (n = 658, AUROC = 0.949). Strong discrimination was also observed in endometrial cancer (n = 422, AUROC = 0.943), bladder cancer (n = 440, AUROC = 0.930), soft tissue sarcoma (n = 417, AUROC = 0.926), renal cell carcinoma (n = 228, AUROC = 0.915), breast cancer (n = 1,682, AUROC = 0.915), non-small cell lung cancer (n = 1,596, AUROC = 0.896), esophagogastric cancer (n = 351, AUROC = 0.889), colorectal cancer (n = 1,064, AUROC = 0.882), melanoma (n = 409, AUROC = 0.850), and ovarian cancer (n = 469, AUROC = 0.845). These results indicate that DeMoP performs well at the cohort level while maintaining broad discriminatory ability across a broad spectrum of solid and hematological malignancies.

Together, these analyses show that DeMoP outperformed the conventional and deep-learning methods evaluated in the GENIE cohort and performed consistently across the tumor lineages examined. The gene-level importance scores highlighted established cancer-associated genes and less well-studied regulators such as ALOX12B, generating testable biological hypotheses for experimental biologists and clinical investigators to follow up.

### DeMoP achieves strong predictive performance in the TCGA cohort

For the TCGA cohort, the initial dataset contained 9,370 samples. Patients were categorized according to overall survival status and duration. Patients who died within 36 months were assigned to the 3-year mortality group, whereas patients who remained alive for more than 36 months were assigned to the long-term survival group. Patients who did not meet either criterion were excluded. After survival-based filtering, 4,470 samples remained. Samples in the 3-year mortality group with all gene-level copy-number values equal to zero were then removed, leaving 4,371 samples. We further excluded patients in the 3-year mortality group for whom both the "*PERSON NEOPLASM CANCER STATUS"* and "*NEW TUMOR EVENT AFTER INITIAL TREATMENT"* columns were missing. The final TCGA cohort consisted of 4,123 samples. We trained DeMoP using 80% of the TCGA cohort and evaluated it on the remaining 20% held-out internal test set. DeMoP achieved the highest class-1 F1 score (0.72) among the benchmarked methods, outperforming logistic regression (0.59), random forest (0.65), XGBoost (0.68), CNN (0.65), LSTM (0.61), and MLP (0.58) (**Fig. 3A**). The pan-cancer setting poses a substantial modeling challenge because tumor types differ markedly in their clinical and molecular profiles. DeMoP addresses this heterogeneity through an MoE architecture that dynamically allocates specialized experts to capture subtype-specific patterns.

**Figure 3.**
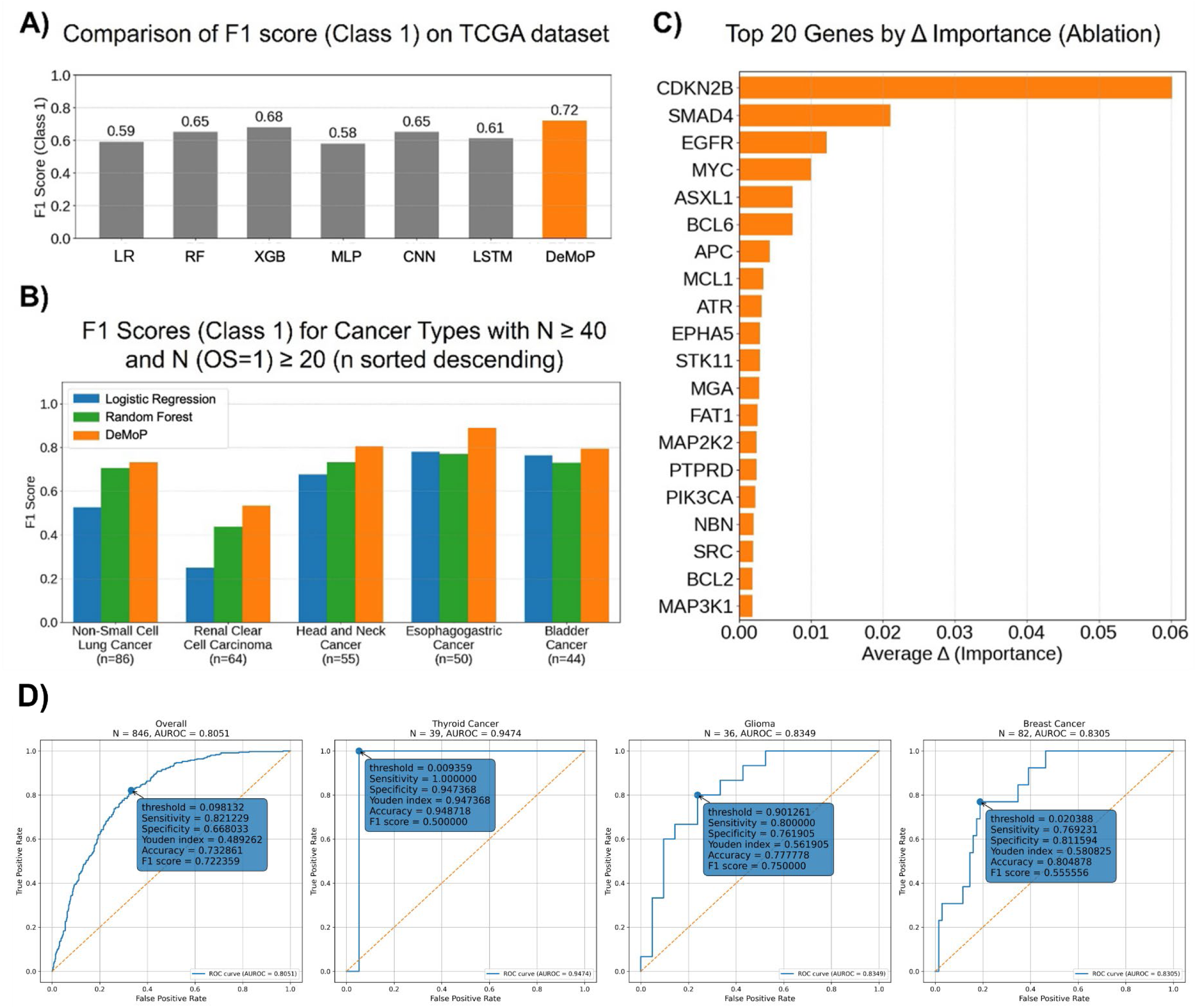
DeMoP performance in the TCGA cohort. (A) Class-1 F1 scores on the held-out test set for DeMoP, logistic regression, random forest, XGBoost, CNN, LSTM, and MLP. (B) Class-1 F1 score comparison of DeMoP, random forest, and logistic regression across tumor types with at least 40 patients and 20 death events (sorted by cohort size). (C) Top 20 genes ranked by the mean signed change in prediction confidence (Δ) in the ablation analysis, with CDKN2B, SMAD4, EGFR, and MYC among the highest-ranked genes. (D) Overall and cancer-specific AUROC performance of DeMoP on the TCGA test set.

We next examined performance by individual cancer type to assess model robustness. **Fig. 3B** shows class-1 F1 scores for five cancer types with at least 40 patients and 20 death events. In non-small cell lung cancer (n = 86), DeMoP achieved a class-1 F1 score of 0.73, compared with 0.70 for random forest and 0.53 for logistic regression. In renal clear cell carcinoma (n = 64), DeMoP achieved a class-1 F1 score of 0.53, more than twice the logistic regression score (0.25) and higher than the random forest score (0.44). DeMoP also outperformed random forest and logistic regression in head and neck cancer (0.81 versus 0.73 and 0.68, respectively), esophagogastric cancer (0.89 versus 0.77 and 0.78), and bladder cancer (0.79 versus 0.73 and 0.76). These results show that DeMoP maintains predictive accuracy even in smaller and more heterogeneous subcohorts, underscoring its potential utility for cancer types where conventional approaches struggle.

Gene-level ablation analysis identified the 20 genes with the highest importance scores (**Fig. 3C**). CDKN2B again ranked highest (importance score = 0.06), indicating consistent model dependence across datasets. SMAD4 (0.02) remained among the top features. Established cancer-associated genes, including EGFR^29^ and MYC^22^ (each 0.01), also ranked highly. Additional genes, such as ASXL1^30^, BCL6^27^, and APC^31^ (≤ 0.01), further illustrate the range of gene-level features identified by the model.

At the cohort level, DeMoP achieved an overall AUROC of 0.805, with sensitivity of 0.791, specificity of 0.684, a Youden index of 0.475, accuracy of 0.729, and a weighted F1 score of 0.712. We further examined cancer-specific discrimination in the test set and identified three cancer types with more than 20 samples and AUROC values above 0.8. Thyroid cancer showed the highest AUROC (n = 39, AUROC = 0.947), followed by glioma (n = 36, AUROC = 0.835) and breast cancer (n = 82, AUROC = 0.831) (**Fig. 3D**). These results show cohort-level discrimination, with AUROC values above 0.8 in the three cancer types examined.

Collectively, these analyses show that DeMoP outperformed the conventional and deep-learning baselines evaluated in the TCGA pan-cancer cohort and achieved higher class-1 F1 scores across the five cancer-specific subcohorts examined. The gene-level importance scores included established cancer-associated genes, supporting interpretation of the model’s gene-level dependencies.

### DeMoP retains predictive utility in cross-cohort transfer

Cross-cohort evaluation is essential for testing whether a model trained in one cohort generalizes to an independent dataset with different sequencing platforms, annotation pipelines, and patient populations. We therefore trained DeMoP on the AACR Project GENIE training subset (n = 50,417) and evaluated it directly on the full filtered TCGA cohort (n = 4,123). The overall class-1 F1 score was calculated using all 4,123 TCGA samples, whereas Fig. 4A also shows cancer-specific results for the eight cancer types with the largest sample sizes. Because GENIE is more than an order of magnitude larger than TCGA, we conducted this one-directional GENIE-to-TCGA transfer to examine whether representations learned from the larger clinical registry retained predictive utility in the independent TCGA cohort.

**Figure 4.**
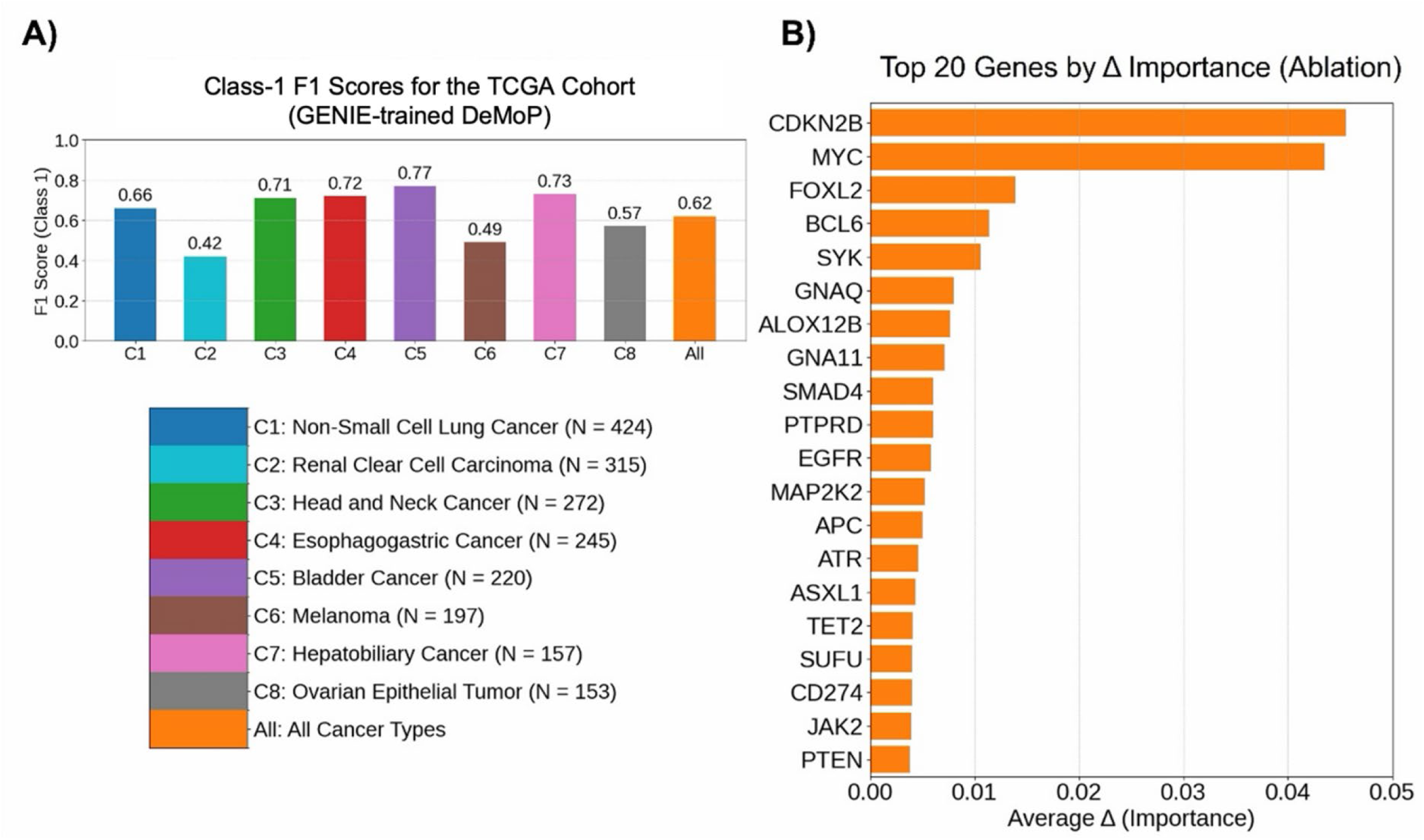
Cross-cohort evaluation of GENIE-trained DeMoP in the TCGA cohort. (A) Class- 1 F1 scores for eight tumor subcohorts (C1–C8), ranging from 0.42 in renal clear cell carcinoma to 0.77 in bladder cancer; overall class-1 F1 = 0.62. (B) Top 20 genes ranked by the mean signed change in prediction confidence (Δ) in the cross-cohort ablation analysis, with CDKN2B (Δ = 0.05), MYC (Δ = 0.04), FOXL2 (Δ = 0.01), and SYK (Δ = 0.01) among the highest-ranked genes.

Transfer performance varied by tumor type. Bladder cancer (C5, n = 220) had the highest class-1 F1 score (0.77), followed by hepatobiliary cancer (C7, n = 157; 0.73), esophagogastric cancer (C4, n = 245; 0.72), and head and neck cancer (C3, n = 272; 0.71). Non-small cell lung cancer (C1, n = 424) had a class-1 F1 score of 0.66, and ovarian epithelial tumor (C8, n = 153) had a score of 0.57. Renal clear cell carcinoma (C2, n = 315) and melanoma (C6, n = 197) had the lowest class-1 F1 scores (0.42 and 0.49, respectively). Across the full TCGA cohort, the transferred model achieved an overall class-1 F1 score of 0.62 (**Fig. 4A**).

Comparison of the cross-cohort results with within-cohort TCGA performance (**Fig. 3A, B**) revealed two patterns. First, the overall class-1 F1 score of 0.62 in the GENIE-to-TCGA transfer setting was lower than the 0.72 observed in the within-cohort TCGA test set, consistent with cohort-specific differences. Second, tumor types that performed well in the within-cohort TCGA analysis, such as bladder cancer (0.79 with the TCGA-trained model; 0.77 with the GENIE-trained model) and esophagogastric cancer (0.89 with the TCGA-trained model; 0.72 with the GENIE- trained model), also performed relatively well in transfer. Renal clear cell carcinoma and melanoma had lower transfer F1 scores. These results indicate that DeMoP retained predictive utility across cohorts, although performance varied by tumor type; application to new clinical registries may benefit from domain adaptation.

In the cross-cohort ablation analysis, CDKN2B again ranked first (importance score = 0.05), indicating consistent model dependence across datasets. MYC (0.04) remained among the top features, consistent with its ranking in the cohort-specific analyses. FOXL2^32^ (0.01) and SYK (0.01) ranked more prominently in the transfer setting. Additional highly ranked genes, including BCL6, GNAQ, and ALOX12B, illustrated both conserved and context-dependent model dependencies.

Comparison of importance scores across the GENIE-only (**Fig. 2C**), TCGA-only (**Fig. 3C**), and GENIE-to-TCGA analyses (**Fig. 4B**) revealed both stable and context-dependent gene-level patterns. CDKN2B and MYC consistently ranked among the top candidate genes across all settings, indicating model dependence on these gene-level features. SMAD4 had higher importance scores in the cohort-specific analyses (0.02 in both GENIE and TCGA) than in the transfer setting (0.006), indicating lower model dependence in the transfer analysis. FOXL2 and SYK also ranked more highly in the transfer setting. Conversely, cancer-associated genes such as PIK3CA^33^ and MAP3K1^34^ ranked lower in the transfer analysis, which may reflect differences in gene-level copy-number alteration prevalence and cohort composition.

In the cross-cohort setting, a model trained on the larger registry retained predictive utility in the smaller independent cohort, although direct transfer remained challenging. These results indicate that representations of clinical and gene-level features transferred across cohorts. Cross-cohort performance may nevertheless benefit from domain adaptation to reconcile platform- and annotation-related differences. Overall, these findings support further evaluation of DeMoP in heterogeneous precision-oncology settings.

## Discussion

DeMoP is a language-model-guided mixture-of-experts framework for cancer prognosis that represents structured clinical variables and molecular data as natural-language sequences and models nonlinear patterns across heterogeneous modalities without manual feature engineering.

Across two independent pan-cancer cohorts, AACR Project GENIE and TCGA, DeMoP outperformed the conventional machine-learning methods and deep-learning architectures evaluated in this study. DeMoP also retained predictive utility when transferred from the larger GENIE registry to the smaller independent TCGA cohort, supporting further evaluation in heterogeneous clinical settings. Beyond predictive performance, gene-level ablation analyses characterized the model’s dependence on individual gene-level segments. The identification of established cancer-associated genes, such as CDKN2B and MYC, and a less characterized candidate, ALOX12B, indicates that predictions depend on both established and less-studied gene- level features. This combination of outcome prediction and gene-level analysis supports the use of DeMoP for prognostic modeling and biological hypothesis generation.

Despite these strengths, several limitations warrant consideration. DeMoP currently integrates clinical variables and copy-number alteration data but does not incorporate additional modalities such as transcriptomics, proteomics, or longitudinal clinical trajectories. Although cross-cohort transfer retained predictive utility, performance varied across tumor types, indicating that domain adaptation or calibration may be needed to address dataset-specific differences. In addition, serializing structured patient features as natural-language sequences may obscure some fine- grained quantitative relationships, motivating hybrid models that combine language-based and structured representations.

Several directions could extend the DeMoP framework. Incorporating additional data modalities, including imaging and single-cell omics, may yield more comprehensive patient representations. Domain-adaptation or continual-learning strategies may improve robustness across institutions and populations. Larger and more diverse training datasets may also help the model capture rare but clinically important patterns. Finally, prospective clinical validation will be necessary to establish the utility of DeMoP as a decision-support tool.

Overall, DeMoP illustrates that a language-model-guided architecture can provide a unified framework for integrating clinical and molecular omics data for cancer prognosis. As foundation models continue to evolve, approaches such as DeMoP may support the analysis of heterogeneous biomedical data for clinical outcome prediction.

## Methods

### Dataset collection and preprocessing

AACR Project GENIE data^19^ were downloaded from Synapse (https://www.synapse.org/Synapse:syn7222066/wiki/405659). TCGA^20^ data were obtained from cBioPortal (https://www.cbioportal.org/). Copy-number alteration data from both TCGA and AACR Project GENIE were available as gene-level annotations and required no additional reformatting. For AACR Project GENIE, overall survival time was defined as the interval between the sample sequencing date and the censoring date. For TCGA, overall survival was obtained directly from the clinical annotations. Patients were then assigned binary outcome labels based on 3-year overall survival. Patients who died within 3 years were labeled class 1, whereas patients who survived for more than 3 years were labeled class 0.

### Text representation

For each patient, we define the prompt string *S*^(*i*)^ as the concatenation of the basic clinical information and the gene-level profile. For example,

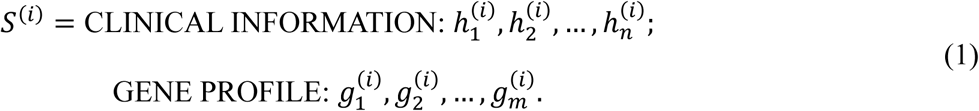

Here, 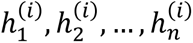 denotes the basic clinical information, and 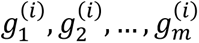 denotes the set of observed gene-level segments, including copy-number alteration data and associated brief descriptive text, for a given patient. This format places numerical and categorical information, together with brief gene descriptions, in a single token sequence, allowing the language model to learn cross-modal dependencies directly from text.

### Patient data encoding

The string *S*^(*i*)^ in equation (1) is tokenized with the DeBERTa-v3 tokenizer18 into a sequence of token indices

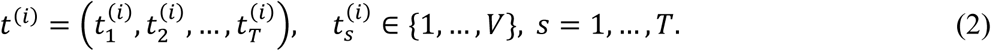

along with an attention mask

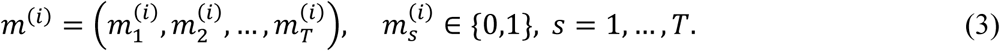

Here, V is the tokenizer vocabulary size, m_t^(i) = 1 indicates a real token, and m_t^(i) = 0 indicates padding. This procedure converts each patient’s mixed-modality record into a uniform sequence of discrete symbols. Each token index t_t^(i) is mapped to a d-dimensional embedding vector e_(t_t^(i)), yielding the input matrix

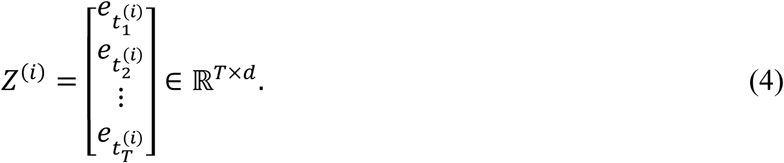

The embedding matrix Z^(i) serves as input to the DeBERTa-v3 transformer encoder, enabling all patient features to be processed through self-attention and feed-forward layers without separate feature-engineering steps.

### DeBERTa-v3-large encoding and layer fusion

We use the pretrained DeBERTa-v3-large encoder as the backbone. Let Z in R^(T x d) be the input embedding matrix for a single patient, where T is the token-sequence length and d = 1,024 is the hidden dimension of each DeBERTa layer. The transformer produces a set of hidden-state tensors

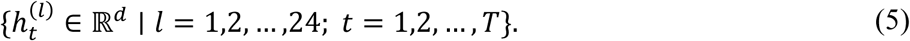

Following the implemented architecture, we concatenate the final four hidden layers at each token position to combine information from the upper contextual layers:

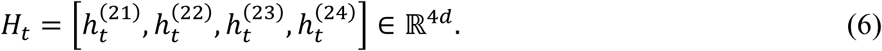

The resulting representation H_t combines multilevel features but has dimension 4d, which is computationally burdensome for downstream modules. To reduce dimensionality while supporting cross-layer interactions, we apply a single-layer MLP with GELU activation35 to each token representation:

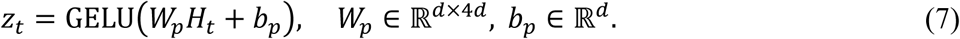

Here, the GELU nonlinearity serves as a learned bottleneck and mixes information from the four layers before pooling. The resulting vector z_t in R^d is a compact fused embedding for token t. The projected sequence {z_t}_{t=1}^T is then passed to the attention-based pooling module and mixture-of-experts head.

### Attention-based token pooling

After projecting each token embedding to z_t in R^d, we aggregate the sequence {z_t}_{t=1}^T into a single vector v in R^d using a lightweight self-attention mechanism9. We first compute query, key, and value matrices by linearly projecting the token embeddings:

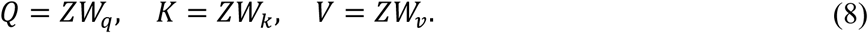

where Z in R^(T x d) stacks the z_t vectors, and W_q, W_k, and W_v in R^(d x d) are learned projection matrices.

We then form the T x T attention weight matrix using scaled dot-product attention followed by softmax normalization:

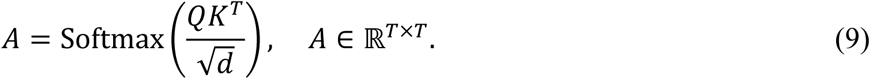

We then compute the context matrix and apply average pooling across tokens:

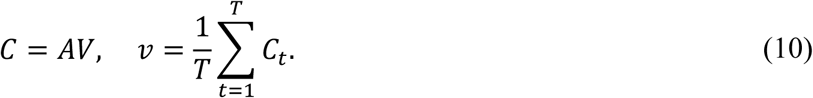

where C_t denotes row t of C. The resulting vector v is a global patient representation that is passed to the mixture-of-experts head for final classification.

### Mixture-of-experts (MoE) head

To model heterogeneous prognostic patterns across patients, we route the pooled representation v in R^d through K parallel expert networks using a soft-gating mechanism. In our implementation, K = 4. The gating module is a two-layer MLP with layer normalization, GELU activation, and dropout:

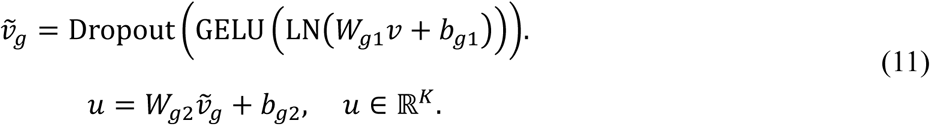

The gating scores are normalized into a distribution over experts:

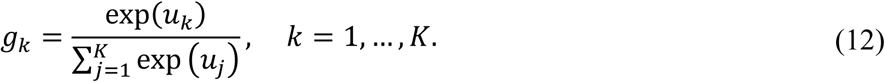

Each expert k applies two sequential ResNet-style fully connected residual blocks to the shared input v. The first block maps the representation from d to h dimensions:

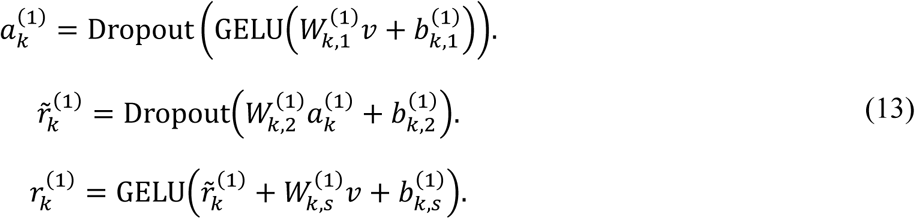

The second residual block maps the representation from h to m dimensions:

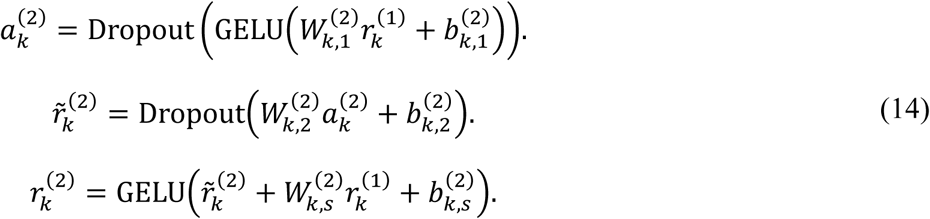

Here, h and m denote the intermediate hidden dimensions, with h = 512 and m = 256 in the implemented model. Shortcut projections are used because the feature dimensions change across residual blocks. Each expert then produces class logits

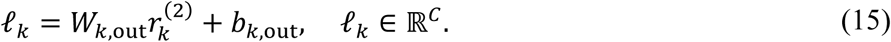

where C = 2 is the number of survival classes. Finally, the gated ensemble combines the expert outputs into the overall logits

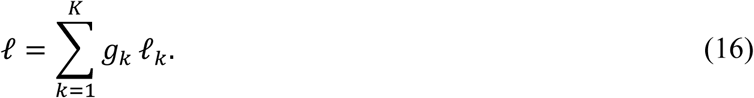

These logits are passed through softmax to obtain class probabilities. The residual shortcuts stabilize gradient flow and encourage each block to learn incremental refinements for survival- group prediction.

### Training objective and optimization

We train the network end to end by minimizing a supervised classification loss. Let y_i denote the ground-truth class label for patient i, and let ell_i,c denote the model logit for class c. The standard cross-entropy loss over N training samples is

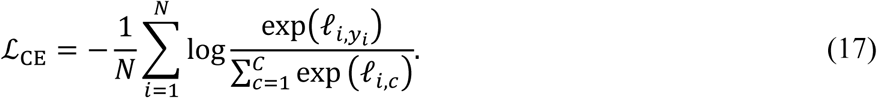

Model parameters are optimized with the AdamW algorithm36, which combines adaptive moment estimation with decoupled weight decay. We use a linear learning-rate schedule with warmup: the learning rate increases linearly from zero to alpha_max over the first W steps and then decays linearly to zero over the remaining training steps. The best checkpoint is selected based on the validation weighted F1 score. In the implemented training callback, early stopping is based on validation loss.

During evaluation, we report accuracy, class-1 and weighted F1 scores, and AUROC. For class c,

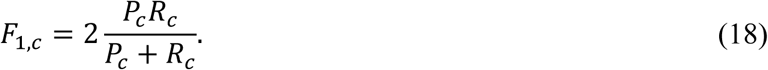

where P_c and R_c denote the precision and recall for class c. The weighted F1 score is computed as

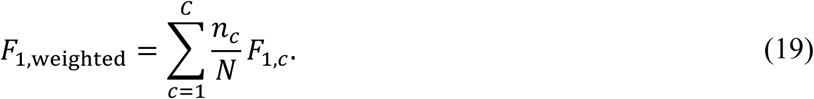

where n_c is the number of samples in class c. This metric accounts for class imbalance by weighting each class-specific F1 score by its class support.

### Ablation study

We performed a systematic gene-level ablation analysis to quantify the dependence of DeMoP predictions on each gene-level segment under the specified input-perturbation scheme. For a given patient *i* and gene *g_j_*, we removed the corresponding gene-level segment, including copy-number alteration information and associated brief descriptive text, when that segment was present. Let *ŷ_i_* denote the class predicted from the original full input (i.e., the class with the highest predicted probability). We then passed the modified input through the fine-tuned model and measured the signed change in predicted probability for the same original predicted class relative to the full input:

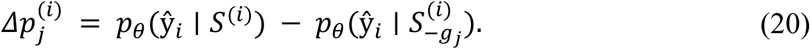

where *p_θ_*(*ŷ_i_* | *S^(i)^*) is the probability assigned to the model’s original predicted class from the full patient description, and *p_θ_*(*ŷ_i_* | *S−_gj_^(i)^*) is the probability assigned to the same class after removal of gene-level segment *g_j_*. The original predicted class is fixed before ablation, even if the predicted class changes after gene removal. Positive values of *Δp_j_^(i)^* indicate that ablation decreases confidence in the original prediction, whereas negative values indicate that ablation increases confidence. We compute the aggregate gene-level importance score as the mean signed change across patients containing that gene-level segment:

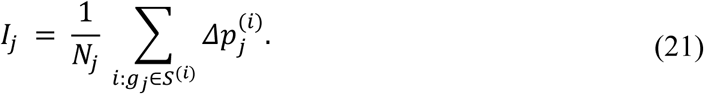

where *N_j_* is the number of patients whose descriptions contain gene-level segment *g_j_*. Genes are ranked by *I_j_* in descending order. Larger positive scores identify gene-level features whose removal, on average, most decreases the model’s confidence in its original prediction. This ranking quantifies model dependence under the specified input-perturbation scheme and does not establish biological mechanism or molecular-driver status.

## Data availability

The datasets used in this study are publicly available, and the original publications for each dataset are cited in the Methods section.

## Code availability

The DeMoP package is available at https://github.com/Lin-Xu-lab/DeMoP.

## Acknowledgments

High-performance computing resources provided by the Quantitative Biomedical Research Center (QBRC) and BioHPC at UT Southwestern Medical Center are gratefully acknowledged.

## Author contributions

CT, LY, and LX conceived and designed the study. CT and LY generated the scripts and GitHub page and performed the data analysis. LX acquired funding. CT, LY, QL, and LX wrote and revised the manuscript. All authors read, revised, and approved the final manuscript.

